# Parameterizing the genetic architecture under stabilizing selection

**DOI:** 10.64898/2026.03.27.714826

**Authors:** Hanbin Lee, Jonathan Terhorst

## Abstract

Across many complex traits, genetic variants with larger effect sizes tend to occur at lower frequencies, which is often interpreted as a signature of stabilizing selection. In statistical genetics, the so-called *α*-model captures this relationship by assuming that effect size variance is inversely proportional to heterozygosity raised to a power 0 ⩽ *α* ⩽ 1. Although empirically useful, the *α*-model is phenomenological rather than mechanistic and lacks a direct population-genetic interpretation. In this paper, we derive an alternative to the *α*-model based on evolutionary theory. Our approach yields a linear mixed model in which the frequency dependence of effect size emerges naturally as a function of interpretable evolutionary quantities describing mutational variance, selection intensity, and coupling between the focal and selected traits. These quantities enter through two identifiable variance components that can be estimated by restricted maximum likelihood (REML). The resulting framework links a fitness-landscape model to standard mixed-model methodology, enabling both inference on evolutionary parameters and downstream prediction by best linear unbiased prediction (BLUP). In forward simulations, the model accurately recovers the focal-trait variance and generally improves genetic prediction relative to conventional *α*-model baselines.

## 1 Introduction

The canonical example of a fitness landscape function is Fisher’s geometric model (FGM), which supposes that an individual’s fitness is given by

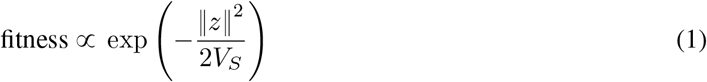

where *z* = (*z*^1^, …, *z*^*d*^) is a *d*-dimensional trait vector subject to stabilizing selection and *V*_*S*_ controls selection intensity (Fisher, 1930; Tenaillon, 2014). Originally proposed by R.A. Fisher almost a century ago, the past decade has seen renewed interest in FGM as a theoretical framework for understanding trait-associated loci in genome-wide association studies (GWAS). By considering a special instance of FGM with isotropically distributed mutational effects, Simons et al. (2018) made predictions about the distribution of (GWAS)-derived effect sizes. A follow-up study used this framework to estimate the contribution of complex traits to fitness in actual GWAS data (Simons et al., 2025). Using predictions derived from FGM, Koch et al. (2024) further argued that stabilizing selection is a primary mode of selection acting on human traits, and gave estimates as to its strength.

These works help to explain repeated empirical observations in massive GWAS of negative correlation between effect size and allele frequency (Zeng et al., 2018; Schoech et al., 2019; Privé, Arbel, and Vilhjálmsson, 2020). Generally, this correlation is ascribed to purifying selection, where large effect variants are more likely to be removed from the population. To model this relationship, a common choice is the so-called *α*-model, by which the variance of the effect size is inversely proportional to allelic heterozygosity raised to the power 0 ⩽ *α* ⩽ 1. This model first appeared in the setting of estimating genetic relatedness matrices from single nucleotide polymorphism (SNP) markers and has later been extended to explicitly detect negative selection (Zeng et al., 2018; Schoech et al., 2019).

Although it captures the underlying relationship well, the *α*-model is largely based on statistical curve-fitting. To the best of our knowledge, a population genetic model that predicts this relationship has yet to be developed (Sella and Barton, 2019; Koch and Sunyaev, 2021). In this work, we fill that gap by proposing a general procedure for combining an evolutionary model with mixed-effects models in statistical genetics. This yields a linear mixed model parameterized by evolutionary quantities that can be estimated through a conventional statistical method, namely the restricted maximum likelihood (REML) estimator (Johnson and Thompson, 1995). These quantities appear in the variance of random effects encoding the aforementioned frequency dependence of effect sizes that is distinct from the *α*-model. Unlike the *α*-model, the new formula does not diverge to infinity as the sample frequency approaches zero and allows unbiased estimation of mutational variance. Furthermore, as a mixed-effects model, it can estimate genetic values from the best linear unbiased predictor (BLUP).

## 2 Theory and model

### 2.1 Diffusion approximation for stabilizing selection

Assuming that individual fitness follows the FGM (1), the marginal selection coefficient *s*_eff_ acting on a given locus is

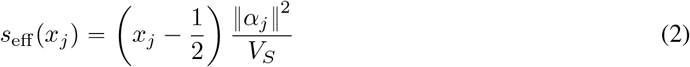

for a biallelic locus *j* of frequency *x* and effect size 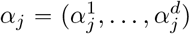 with 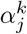 corresponding to *z*^*k*^ (Robertson, 1956; Lande, 1976; Simons et al., 2018; Negm and Veller, 2026). Hereafter, *α*_*j*_ denotes the vector of effect sizes of the *j*-th locus for traits *k =* 1, …, *d* and 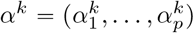 denotes the vector of effect sizes of the *k*-th trait for loci *j =* 1, …, *p*.

In a panmictic diploid population of size *N*, the diffusion approximation corresponding to (2) is

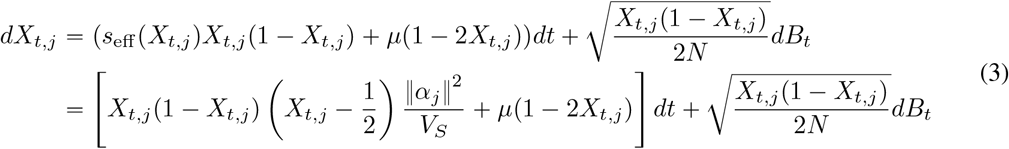

where *B*_*t*_ is the Brownian motion. Rescaling time in units of 2*N* generations, with *V*_*S*_ *=* 2*N* = *W*_*S*_ and *θ =* 4*Nµ*, we obtain

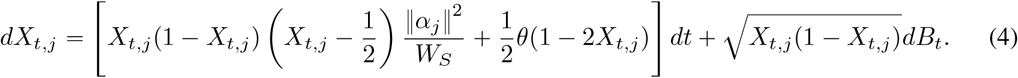

The forward Kolmogorov equation corresponding to (4) is

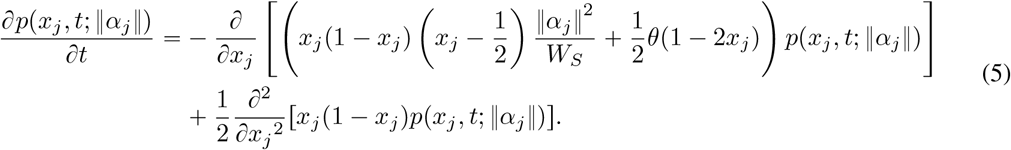

At stationarity, ∂*p*/∂*t =* 0 and we obtain (Ewens, 2004)

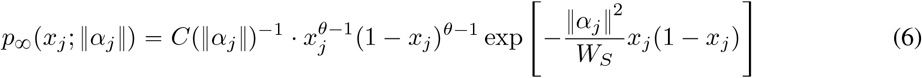

with normalizing constant

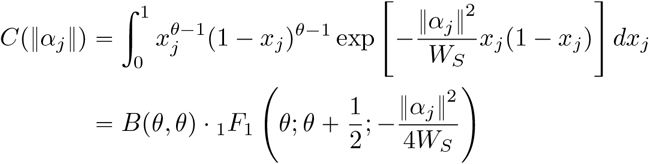

where _1_*F*_1_(*a*; *b*; *z*) is the Kummer’s hypergeometric function (Standards and (U.S.), 2010). For fixed *b, z* we have _1_*F*_1_(*a*; *b*; *z*) = 1 + 𝒪 (*a*). Hence, as *θ* → 0,

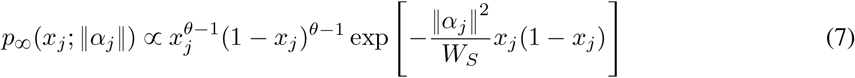

after suppressing *C*(‖*α*_*j*_‖) ≈ *B*(*θ, θ*) which is independent of both *α*_*j*_ and *θ*.

### 2.2 Connecting evolution to mixed-effects models

Genomic mixed-effects models assume the conditional distribution of *y* given *G* is

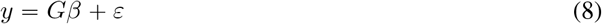

where *y* is the focal trait and *β* is the corresponding effect size. Accordingly, the variance decomposition of the trait is

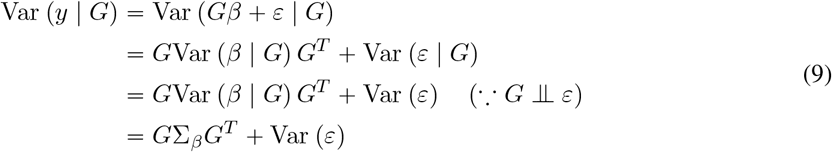

where Σ_*β*_ *=* Var (*β* |*G*).

The aforementioned *α*-model supposes that

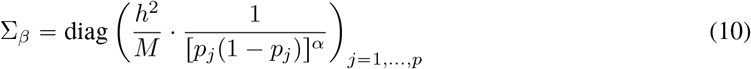

where *M* is the number of variants and *h*^2^ is a positive constant often referred to as the single nucleotide polymorphism (SNP) heritability. Here, the effect of *G* enters the formula through its column-wise frequency *p*_*j*_. In practice, we only have the sample frequency 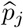 instead of *p*_*j*_, leading to the plug-in estimator

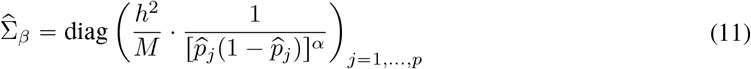

replacing Σ_*β*_. The *α* = 1 case is the genome-wide complex trait analysis (GCTA) model (Yang et al., 2011). Together with *G*, 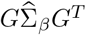 is a genomic estimate of genetic relatedness, called the genomic genetic relatedness matrix (GRM) (Speed and Balding, 2014). For a more comprehensive review of different choices of Σ_*β*_, see Lehmann et al. (2025) and Weir and Goudet (2026).

Despite the success of equation (11) in data modeling, there are some challenges relating to theory and interpretation. First, the form of Σ_*β*_ *=* Var (*β*|*G*) in equation (10) is simply assumed *a priori*, so the parameters *h*^2^ and *α* are merely tuning parameters without an explicit biological interpretation. *α* in particular exists in order to heuristically capture the inverse relationship between minor allele frequency and effect size (Zeng et al., 2018; Schoech et al., 2019). For *α* > 0 there is also the pathology that [*p*_*j*_(1 − *p*_*j*_)]^−*α*^ → ∞ as *p*_*j*_ → 0, requiring heuristic GRM partitioning over multiple MAF bins if rare variants are involved (Yang et al., 2015; Wainschtein et al., 2022). Finally, it is unclear if the sample frequency 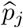 can replace the true frequency *p*_*j*_, especially for rare variants that tend to exhibit larger sampling variance.

Our framework offers a principled solution to the aforementioned issues. The key is to *deduce*, instead of assuming, a formula for Σ_*β,j*_ = Var (*β*_*j*_ | *G*) from evolutionary theory where *j* denotes the *j*-th locus. A common assumption in the literature is that the mean of *β*_*j*_ is zero, thereby we get 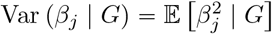.

Then, we can derive 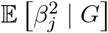 from

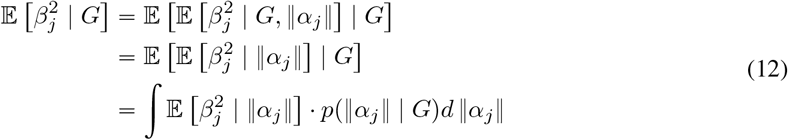

where *α*_*j*_ is the effect size *vector* of the *j*-th locus (previously we used the superscript *k* on *α* to denote the corresponding *k*-th trait).

### 2.3 Fitness effect and its influence on a focal trait

From (12), it remains to determine 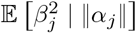, which measures how much the focal trait’s effect *β*_*j*_ is determined by the overall fitness effect *α*_*j*_, and *p*(‖*α*_*j*_‖ |*G*), the likely value of overall fitness given the genotype data from a given evolutionary model. We now derive these quantities in the case of stabilizing selection.

For the rest of the paper, we assume

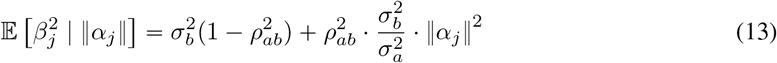

for some constants 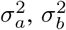 and 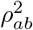. This can be viewed as a generalization of the framework proposed by Simons et al. (2018): with an isotropic FGM,

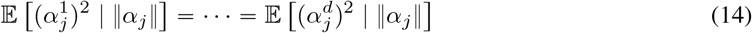

holds by symmetry (all 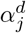 have the same expected magnitude) and

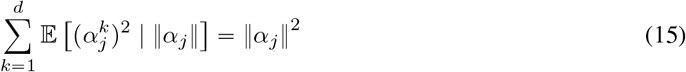

gives 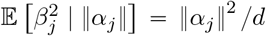 where *β*_*j*_ is one of the 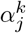. Simons et al.’s model is recovered by setting 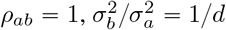.

Another scenario that leads to (13) is when the fitness effect is drawn from a multivariate normal distribution

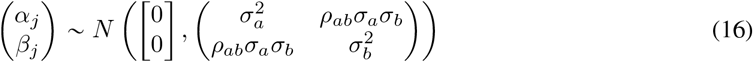

and selection is acting on a one-dimensional trait with fitness effect *α*. Then, 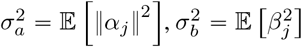, and 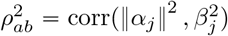 from the property of multivariate normal distributions.

### 2.4 Approximate genetic architecture

Next we turn to *p*(‖ *α*_*j*_‖ ^2^ | *G*). Assuming linkage equilibrium, we have

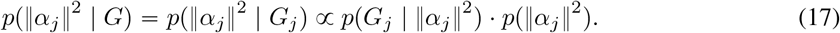

For the first term on the right, *p*(*G*_*j*_| ‖ *α*_*j*_‖^2^, assume that there are *k*_*j*_ derived alleles observed in *G*_*j*_. By exchangeability of the 2*n* samples,

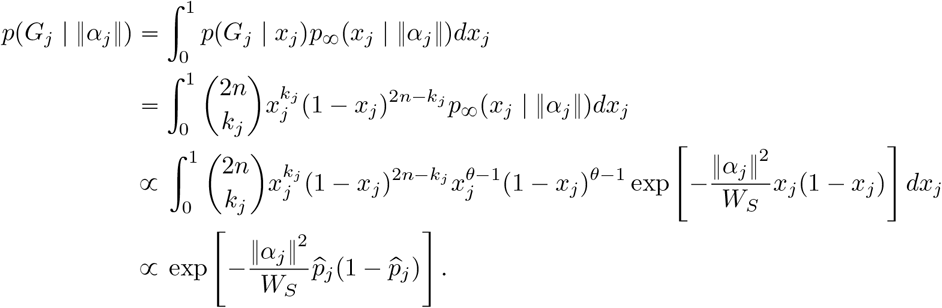

Note that the integration over [0, 1] collapses to the substitution 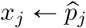 because the beta-binomial distribution degenerates to the sample frequency in large samples (*n →* ∞). For the prior term *p*(‖*α*_*j*_‖^2^), we assume ‖ *α*_*j*_‖ follows a *χ*^2^-distribution multiplied by 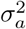. Then 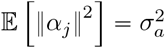 and

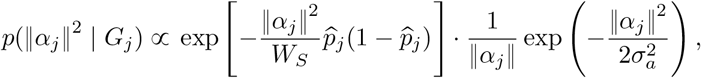

Yielding

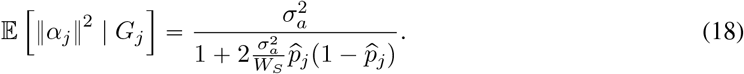

Substituting (18) and (19) into (12), we obtain

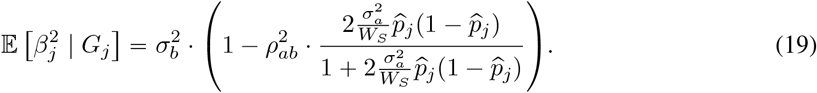

## 3 Simulation and empirical study

### 3.1 SLiM forward simulation

To evaluate our theoretical model, we conducted forward simulations using SLiM (Haller, Ralph, and Messer, 2025). Our individual-based simulations assume that stabilizing selection acts according to the fitness function in equation (1). The simulation is based on (16), where a one-dimensional trait is subject to selection while the observed trait is correlated with it through its focal effect sizes (precise details are given in Appendix B). Notably, this simulation setup differs from most other LMM benchmarking studies, which typically condition on fixed genotypes and only vary the random effects and the residual noise of the regression model (Speed et al., 2017; Gazal et al., 2019; Privé, Arbel, and Vilhjálmsson, 2020).

### 3.2 Variance component estimation

We evaluated equation (19) by simulating traits with small non-genetic noise 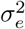. For each replicate, we fit the evolutionary mixed model to the full genotype matrix after removing monomorphic sites and computing the sample allele frequencies 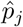 from the remaining dosage columns. These frequencies were inserted into the variance formula in equation (19), and the phenotype was analyzed after adding Gaussian noise of fixed variance 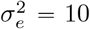. With *ρ*_*ab*_ fixed to its simulation value, we then maximized the marginal likelihood over the two log-transformed parameters 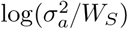 and the overall prior scale parameter 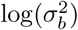 using an L-BFGS-B optimizer implemented in `scipy` (Virtanen et al., 2020).

We first examined whether REML recovers the variance components implied by equation (19). Figure 1 shows that the fitted values preserve the ordering of the true parameter combinations across the simulation grid, but they lie systematically below the naive reference 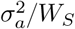. The discrepancy is most visible when selection-induced frequency dependence is strongest (larger 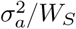). At the same time, the estimates are substantially closer to the Bulmer-corrected reference lines than to the uncorrected target, indicating that the main departure from the single-locus theory is consistent with attenuation from linkage disequilibrium rather than a failure of the functional form itself. See Appendix A for additional details about the correction. The 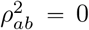 panels are uninformative, because selection on the latent trait does not induce frequency dependence in the focal trait when the two traits are uncorrelated. Formally, one can see in equation (19) that the frequency-dependent term entirely vanishes when 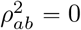.

**Figure 1.**
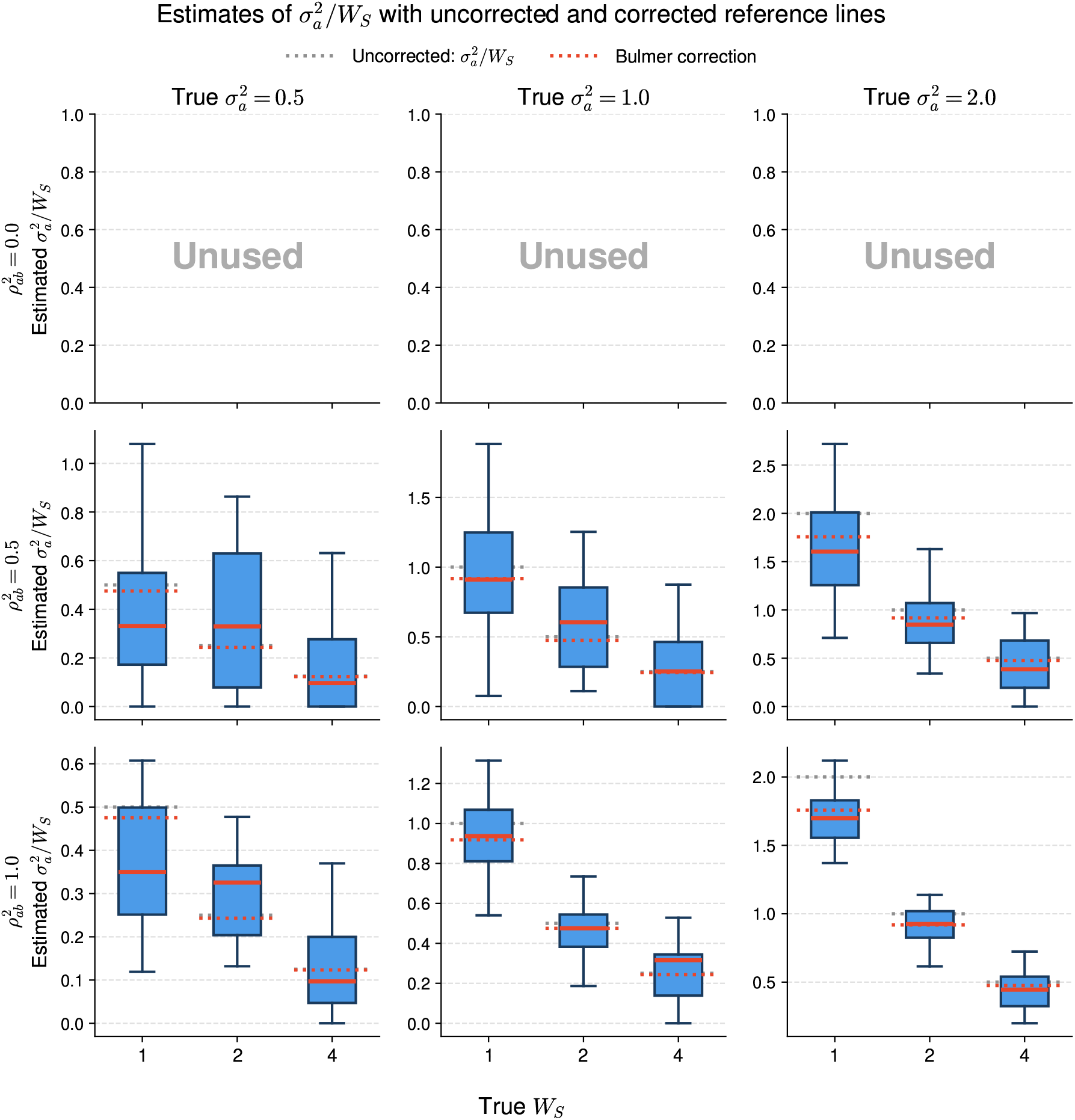
REML estimates of 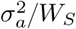 across forward-simulation replicates. Columns correspond to the true 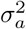, rows correspond to the true 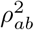, and the x-axis gives the true *W*_*S*_. Gray dotted lines show the uncorrected target 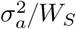, while red dotted lines show the Bulmer-corrected reference derived in the appendix. The 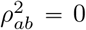 panels are unused because the focal trait then carries no information about selection on the latent trait.

The overall scale parameter 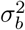 is considerably easier to recover. Figure 2 shows that REML estimates of 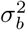 remain centered near the true value across the full grid of *W*_*S*_ and 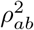, with only modest replicate-to-replicate variability. This further supports the robustness of the functional form of equation (19). Taken together, these results suggest that the focal-trait variance is well identified, whereas the selection-sensitive component is recovered chiefly through the combined quantity 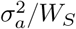 and is therefore more sensitive to departures from the single-locus approximation.

**Figure 2.**
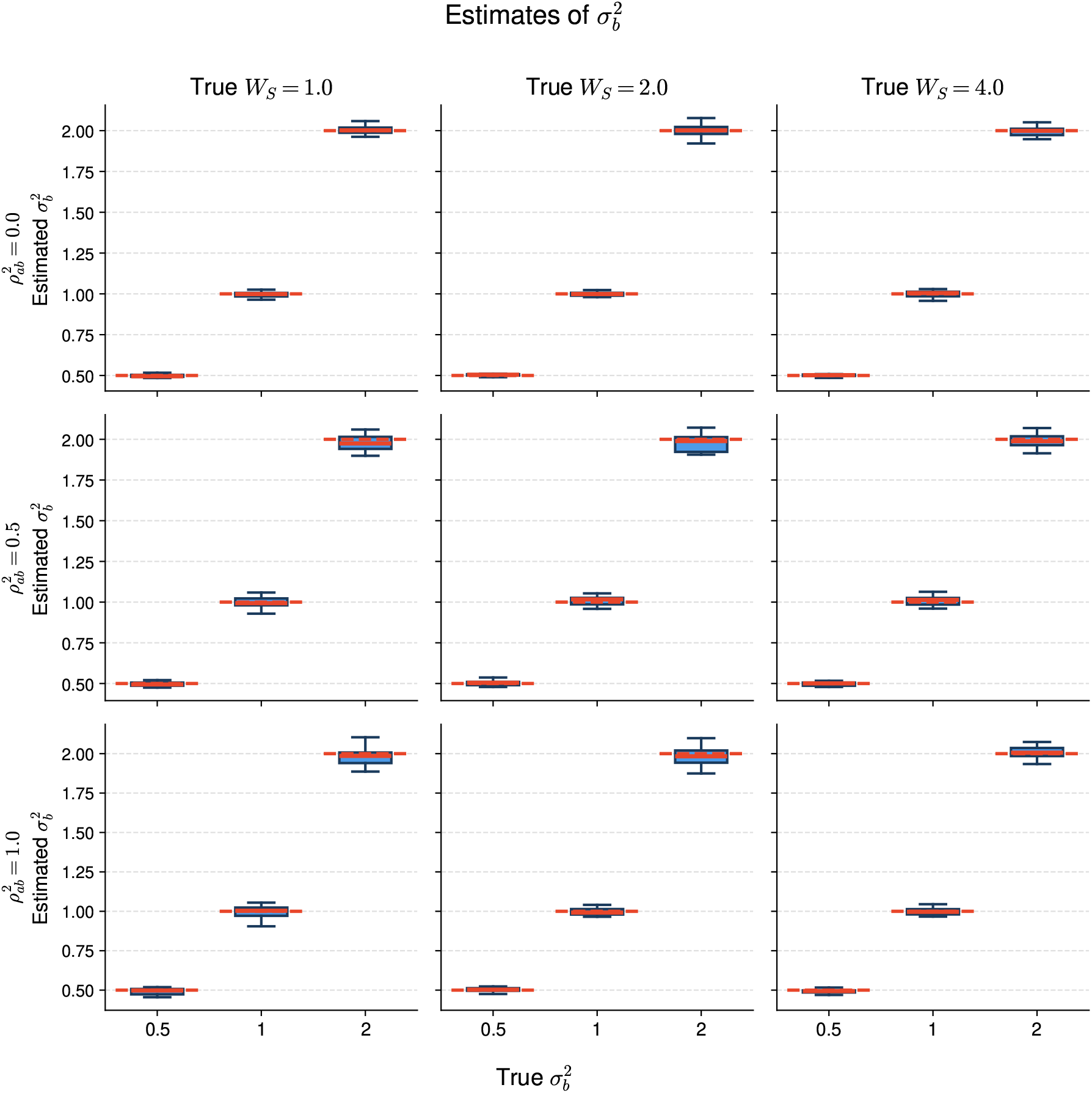
REML estimates of 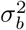 across forward-simulation replicates. Columns correspond to the true *W*_*S*_, rows correspond to the true 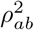, and the x-axis gives the true 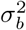. Horizontal reference lines indicate the true values.

### 3.3 Genetic prediction

For the prediction experiments, we first added Gaussian noise of variance 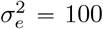 to each replicate phenotype, then split individuals into equally sized training and validation sets using the replicate identifier as the random seed. All fits were carried out on the training set after centering by the training mean. Under the evolutionary model, we jointly estimated 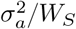, the overall variance scale 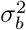, and 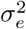 by REML. For the *α*-model baselines with *α* ∈ {0, 0.5, 1}, we instead fit the variance scale and 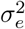 by REML using per-variant prior variances proportional to 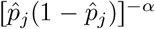. The fitted prior variances were then used to form posterior mean BLUP effect estimates on the training set, which were evaluated on the held-out individuals by *R*^2^.

We next compared held-out BLUP accuracy under the evolutionary model and three *α*-model baselines. Figure 3 reports the validation *R*^2^ across the same grid of simulation parameters. The strongest pattern is the poor performance of the *α* 1 model, which underperforms substantially in every regime. The *α* 0.5 model is closer, but still remains below the evolutionary fit throughout most of the grid. By contrast, the evolutionary model and the *α* “ 0 model are often quite similar, especially when 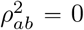, where equation 19 reduces to an approximately frequency-independent architecture. When the coupling between the focal and selected traits is stronger, the evolutionary model typically retains a nearly unnoticeable advantage.

**Figure 3.**
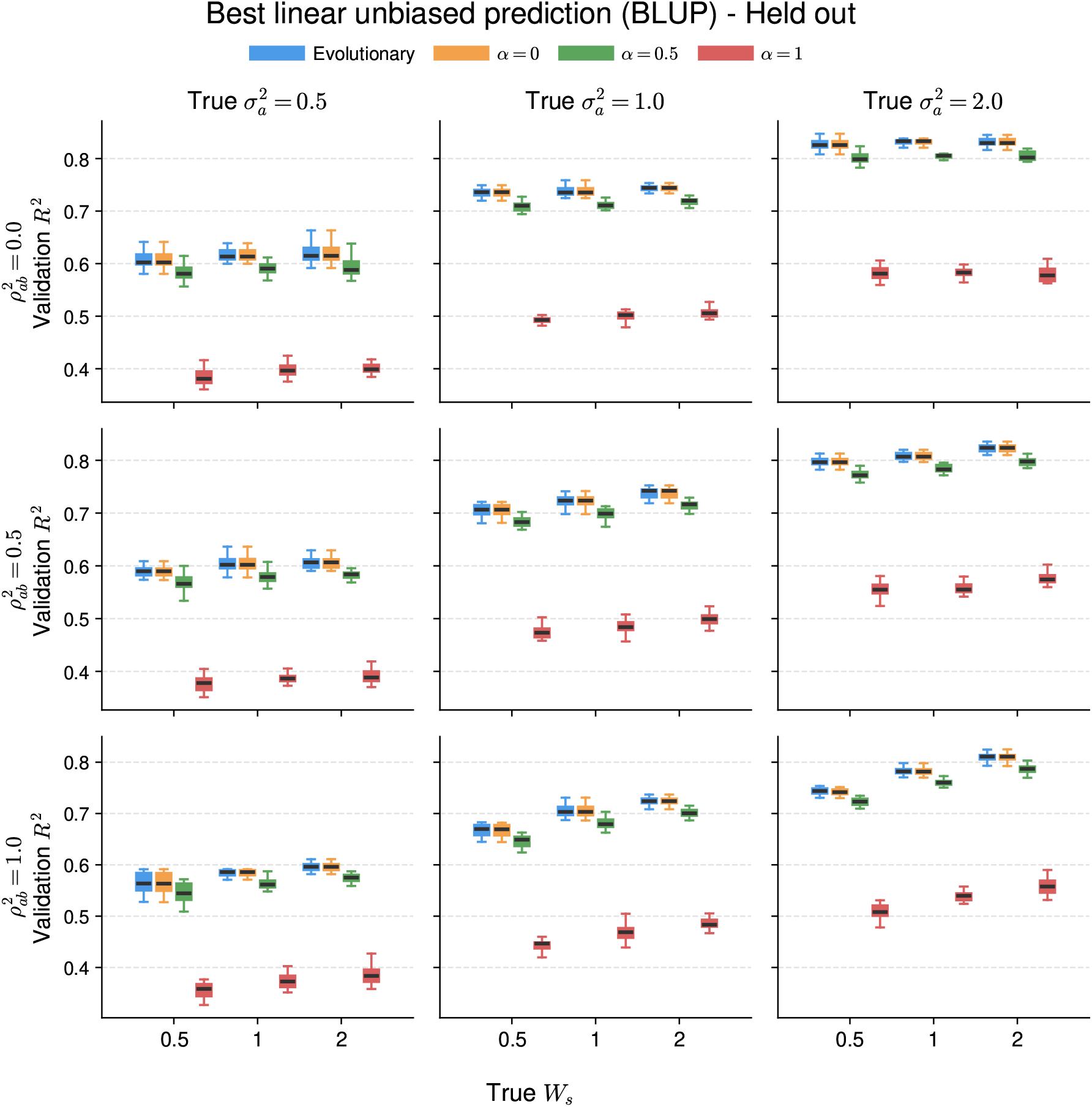
Held-out BLUP validation *R*^2^ for the evolutionary model and the *α*-model baselines. Columns correspond to the true 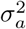, rows correspond to the true 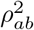, and the x-axis gives the true *W*_*S*_. Error bars summarize variation across replicate populations.

The distinction is sharper when one examines the variance components obtained from the prediction fits. In Figure 4, the evolutionary model remains close to the true 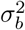 throughout the simulation grid. The *α* = 0 model is approximately unbiased only when 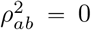 and becomes downward biased once the focal trait inherits frequency dependence from the selected trait. The degradation is more pronounced in smaller values of *W*_*S*_ corresponding to stronger selection. The *α =* 0.5 and *α =* 1 fits underestimate 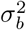 much more severely in all scenarios. Hence, the similarity in predictive accuracy between the evolutionary model and *α =* 0 in some panels should not be interpreted as recovery of the same architecture: the heuristic model can predict reasonably well while still mischaracterizing the underlying variance components. Overall, this shows that the *α*-model fails to capture the frequency-dependent genetic architecture properly.

**Figure 4.**
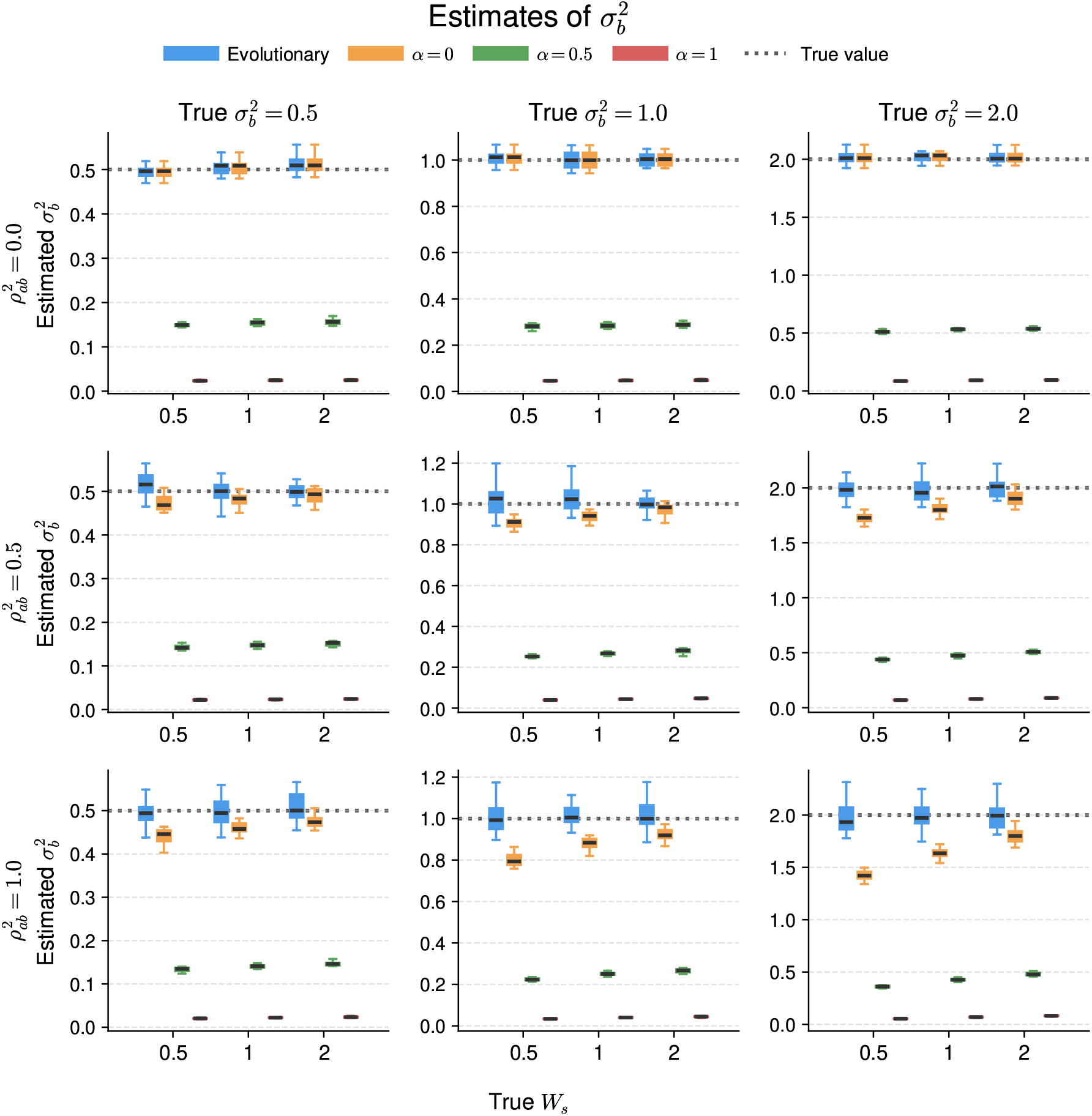
Estimates of 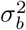 obtained from the BLUP fits. Columns correspond to the true 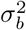, rows correspond to the true 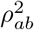, and the x-axis gives the true *W*_*S*_. The dotted horizontal line marks the true value.

Figures A1 and A2 repeat the same training/validation pipeline but fix the evolutionary fit at 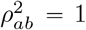. Held-out prediction is nearly unchanged relative to Figure 3, and estimation of 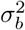 remains close to the full model, although with somewhat greater dispersion in some settings. This supports the reduced parameterization as a useful approximation when separate identification of 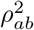 and 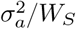 is weak.

## 4 Discussion

We presented a general procedure for deriving evolution-aware mixed-effects models and applied it to stabilizing selection with pleiotropy. This yields a frequency-dependent effect-size formula expressed directly in terms of evolutionary parameters, while still allowing those parameters to be estimated by restricted maximum likelihood (REML). In that sense, the framework links an interpretable fitness-landscape model to standard statistical-genetic practice: the same model can be used both to learn about the evolutionary architecture of a trait and to perform downstream tasks such as genetic prediction by best linear unbiased prediction (BLUP). In forward simulations, this evolutionary model outperformed the conventional *α*-model across the values of *α* we considered, which supports the practical value of deriving the frequency dependence from an explicit evolutionary model rather than imposing it heuristically.

A central ingredient in this construction is the model of pleiotropy. Empirical studies suggest that pleiotropy is pervasive in human complex traits, and Simons et al. (2018) captured this idea within Fisher’s geometric model (FGM) by assuming isotropic mutational effects. Under that assumption, each axis of the underlying fitness landscape contributes equally, so the effect of a mutation on the focal trait is spread symmetrically across directions in trait space. Equation (13) generalizes this relationship by allowing the focal trait to be coupled to overall fitness in a potentially non-isotropic way. One concrete example is a jointly Gaussian model for mutational effects, which has been studied in earlier theoretical work and which we also used in our simulations (Kimura and Crow, 1964; Bürger, 1986; Bürger, 2000; Walsh and Lynch, 2018). More broadly, this suggests that the present framework should not be tied to a single pleiotropy mechanism: an important next step is to investigate what other biological models lead to the same or similar relationship in equation (13).

The simulations also clarify what aspects of the model are practically identifiable. In particular, jointly estimating *ρ*_*ab*_ and 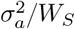 is difficult because 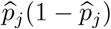 is typically both small and weakly variable under the *U* -shaped allele-frequency spectrum. As a result, the denominator in equation (19) is usually close to one, so the data mainly identify the product 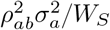 rather than its two components separately. This is why fixing *ρ*_*ab*_ = 1 and interpreting the fitted quantity as 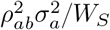 works reasonably well in practice. As we show in Appendix A, this reduced parameterization lowers precision somewhat but has little effect on BLUP accuracy, and estimation of 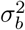 remains stable. That asymmetry is itself informative: 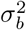 controls the overall scale of the covariance structure, whereas 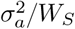 enters only through a weaker modulation by allele frequency. Even when the evolutionary decomposition is only partially identified, the model can therefore still recover the aggregate variance carried by the focal trait.

This perspective also helps explain the behavior of the *α*-model baselines. When 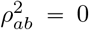, the focal trait is effectively decoupled from the selected background trait, so stabilizing selection induces essentially no frequency dependence and the model collapses to an approximately frequency-independent prior. That regime is therefore a useful sanity check: the corresponding panels are uninformative for 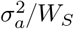, and the evolutionary fit and the *α* = 0 baseline give nearly identical predictions. More generally, because most variants are rare, one often has 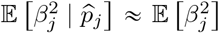, so the genetic relatedness matrices produced by the evolutionary model and the *α =* 0 model differ mainly by an overall scale factor to which BLUP is largely insensitive. Nevertheless, that similarity in prediction should not be mistaken for recovery of the same architecture. Once the focal trait is coupled to the selected background, the *α =* 0 model absorbs the missing frequency dependence by pushing its estimate of 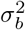 downward, whereas the nonzero *α*-models degrade more substantially in both prediction and variance-component estimation. The comparison therefore shows that predictive performance alone can obscure architectural misspecification, and that an evolutionarily derived frequency-dependence model is preferable to purely statistical heuristics.

Finally, these results should be interpreted in light of the simplifying assumptions used to obtain a tractable formula. Our derivation relies on genotype-likelihood factorization in the spirit of composite likelihood, stationarity of the population, a low mutation rate, and weak selection relative to recombination. In the full multilocus setting, the dynamics of a focal locus depend on the genetic background through linkage, even in the strictly additive setting considered here, and ignoring that dependence lets us reduce the analysis to a single locus. We suspect that this omission is responsible for the systematic underestimation of 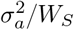 seen in the simulations. In particular, linkage disequilibrium induces the Bulmer effect, which creates background forces opposing selection and thereby weakens the effective selective pressure acting on a locus. Negm and Veller (2026) formalized this reduction as a multiplicative factor, which algebraically has the same effect as increasing the selection width *W*_*S*_ and hence decreasing 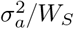. By expanding their formula with further approximations, we found that the Bulmer effect quantitatively predicts the observed underestimation; Appendix A gives the details. Extending the framework to treat linkage disequilibrium and richer models of pleiotropy more explicitly is therefore the most natural direction for future work.

## Acknowledgements

This work greatly benefited from discussions with Josh Schraiber (University of Southern California) and Guy Sella (Columbia University). H.L. thanks the attendees of New York Area Population Genetics 2026 for valuable comments. This research was supported in part by the National Institute of General Medical Sciences of the NIH under award number R35GM151145. The content is solely the responsibility of the authors and does not necessarily represent the official views of the NIH.

## A Mean-field approach to linkage disequilibrium

So far, we completely ignored the effect of linkage which creates theoretical challenges. To analyze the dynamics of an individual locus in general, we have to consider the countless number of interactions with other loci that exhibit randomness. To simplify the problem, we adopt the *mean-field* approximation where the locus is assumed to interact with the rest of the system in a simplified fashion. Here, the field is the rest of the genome, or the genetic background, with which the focal locus interacts. Because this background consists of a huge number of loci, although being stochastic in principle, we instead treat it as a deterministic *average* which gives rise to the term mean-field.

We work with each locus while leaving all else as a deterministic aggregate. Under this framework, one can further refine equation (2) as (after scaling by 2*N*)

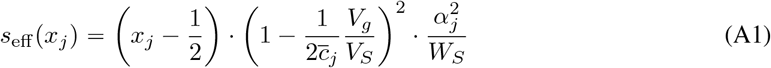

where 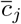 and 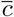 are local and global harmonic means of the recombination fractions, respectively, given by

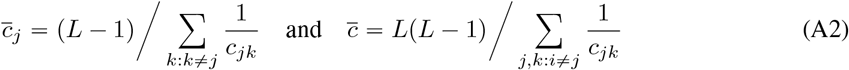

and 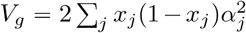 is the genic variance of *z* as derived in Negm and Veller (2026). For simplicity, let *u*_*j*_ be the normalized position of locus *j* so that *Lu*_*j*_ is the position in base pairs, and write 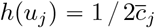.

The difficulty arising in equation (A1) is that *V*_*g*_ is also a random variable because it is a function of the random variables *x*_*j*_. We take another round of mean-field approximation so that *V*_*g*_ *=* 2*L* · 𝔼 [*x* (1 *x* − *α*^2^)] is a constant. By further assuming that mutation rate is uniform, otherwise we can rescale the genome, we arrive at

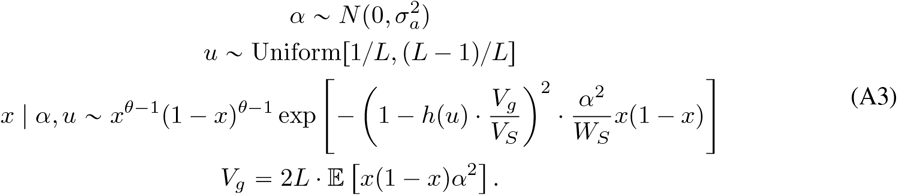

under stationarity as before. Solving for *V*_*g*_ with *h*(*u*)*V*_*g*_ ≪ *V*_*S*_ gives

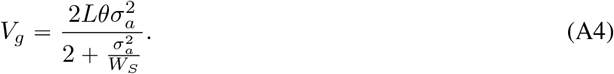

which means that the shape of *h* is irrelevant under the *V*_*g*_ ≪ *V*_*S*_ approximation. A locus-level equivalent is derived in Keightley and Hill (1988). With diminishing selection *W*_*S*_ → ∞, the formula reduces to the classic neutral result 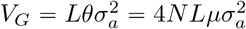 (Lynch and Hill, 1986).

We now follow the same steps as before to deduce

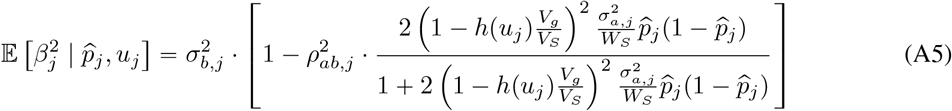

which reduces to

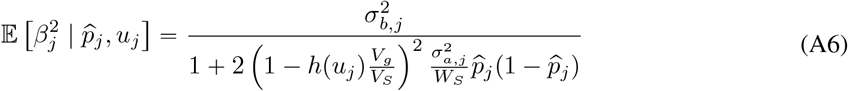

if *ρ* = 1, a perfect correlation between *α* and *β*. As before, if *ρ* = 0, it simply becomes 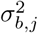 because selection on *z* has no influence on *y* at all. For the correction factors in Figure 1, we used the global harmonic mean of recombination fraction.

## B Simulation details

The population followed a panmictic demographic history with a constant diploid population size of *N =* 10,000 individuals. Each genome was of length *L =* 2 × 10^6^ with mutation rate of *µ =* 10^−8^ and recombination rate of *r =* 10^−5^. The configuration was borrowed from Hayward and Sella (2022) to set *Lµ =* 0.02 mutations per genome per generation, 2*NLµ =* 400 ≫ 1 to ensure polygenicity, and *r/µ =* 10^3^ for low linkage disequilibrium (see Appendix A). The mutational effect sizes *α* were simulated from 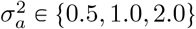 and the selection width *W*_*S*_ *= V*_*S*_*/*2*N =* {1.0, 2.0, 4.0} (see Appendix for details). As *V*_*S*_ is of scale 𝒪(*N*), the environmental variance *V*_*E*_ and the genetic variance *V*_*G*_ were negligible. Hence, the fitness function (1) was used in the simulations. For each pair of 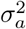 and *W*_*S*_, a single burn-in of 10*N* generations was followed by 20 separate replicates of 5*N* generations.

To simulate pleiotropic selection, for each population after a total of 15*N* generations, we drew the observed effect sizes *β* conditional on *α* according to the joint distribution in equation (16). This is the same as sampling *β* during the SLiM simulation because *β* has no effect on the simulation at all (*α* does the job). For 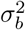 and 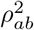, we chose them to be the same as 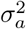 and to take values in the grid {0.00, 0.50, 1.00}, respectively.

**Figure A1:**
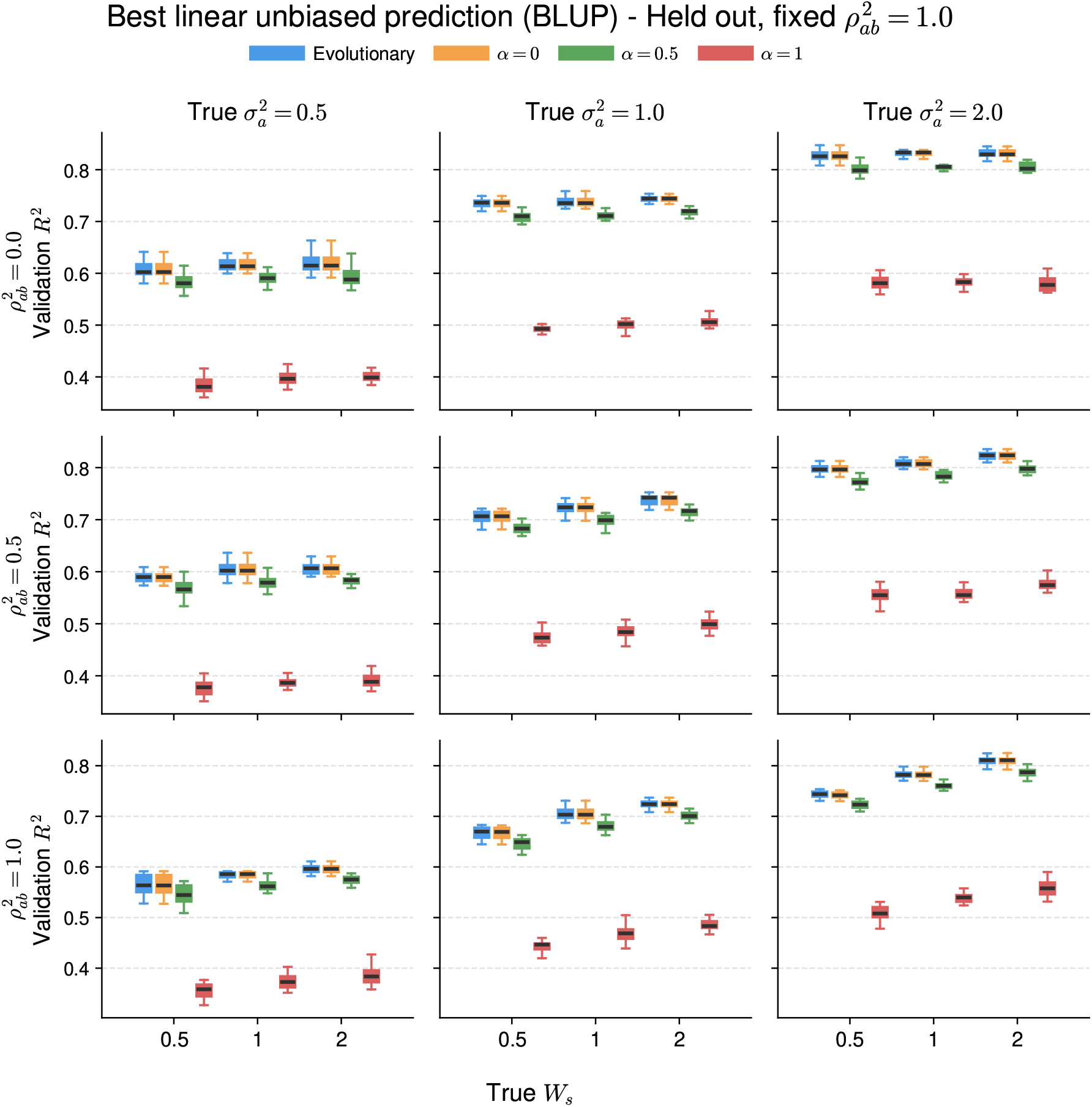
Held-out BLUP validation *R*^2^ when the evolutionary model is fit with 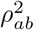 fixed to 1. Columns correspond to the true 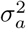, rows correspond to the true 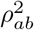, and the x-axis gives the true *W*_*S*_. Error bars summarize variation across replicate populations.

**Figure A2:**
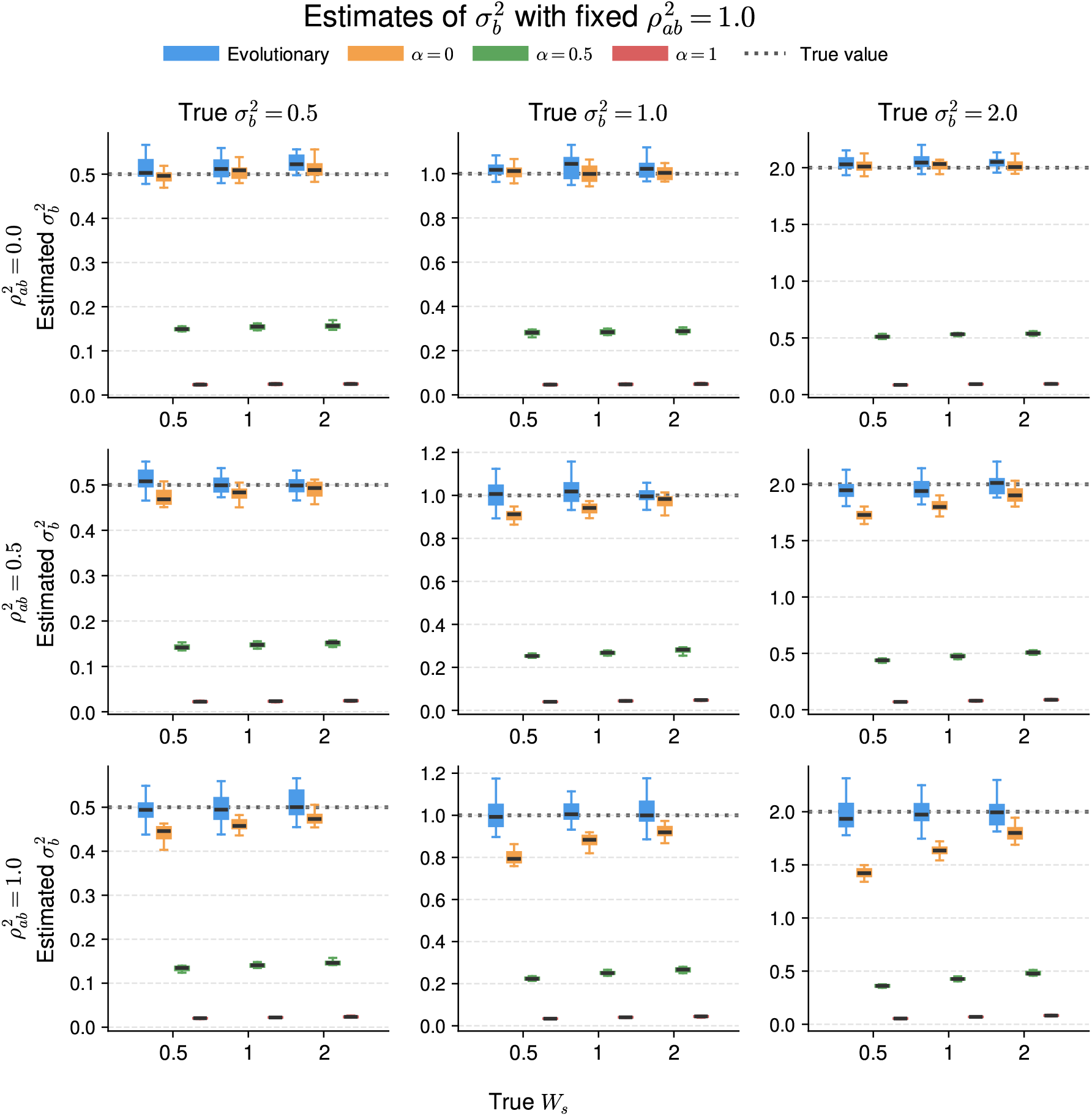
Estimates of 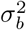 when the evolutionary model is fit with 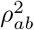 fixed to 1. Columns correspond to the true 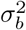, rows correspond to the true 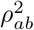, and the x-axis gives the true *W*_*S*_. The dotted horizontal line marks the true value.

